# DomeVR: A setup for experimental control of an immersive dome virtual environment created with Unreal Engine 4

**DOI:** 10.1101/2022.04.04.486889

**Authors:** Katharine A. Shapcott, Marvin Weigand, Iuliia Glukhova, Martha N. Havenith, Marieke L. Schölvinck

## Abstract

Immersive virtual reality (VR) environments are a powerful tool to explore cognitive processes ranging from memory and navigation to visual processing and decision making - and to do so in a naturalistic yet controlled setting. As such, they have been employed across different species, and by a diverse range of research groups. Unfortunately, designing and implementing behavioural tasks in such environments often proves complicated. To tackle this challenge, we created DomeVR, an immersive VR environment built using Unreal Engine 4 (UE4). UE4 is a powerful game engine with photo-realistic graphics containing a visual scripting language designed for use by non-programmers. As a result, virtual environments are easily created using drag-and-drop elements. DomeVR aims to make these features accessible to neuroscience experiments. This includes a logging and synchronization system to solve timing uncertainties inherent in UE4; an interactive GUI for scientists to observe subjects during experiments and adjust task parameters on the fly, and a dome projection system for full task immersion in non-human subjects. These key features are modular and can easily be added individually into other UE4 projects. Finally, we present proof-of-principle data highlighting the functionality of DomeVR in three different species: human, macaque and mouse.

## 2 Introduction

In recent years it has become abundantly clear that the study of brain activity will benefit enormously from immersive naturalistic tasks [1, 2, 3, 4, 5]. Traditionally, neuroscience has taken a reductionist approach: rigorous experiments with simplified stimuli were designed to dissect out single cognitive processes, such as attention [6] or memory [7], and their neuronal underpinnings. Even though this approach has been immensely successful in explaining brain activity acquired during such experiments, it is hardly reflective of the highly dynamic and varied environment that the brain encounters in everyday life. Indeed, the use of more naturalistic visual input has forced us to revise such fundamental notions about the brain as the receptive field [8, 9, 10, 11]. However, truly naturalistic settings (i.e. animals freely roaming) limit experimental control, such as precise tracking of eye movements and the exact timings of events to relate the neural activity to. To circumvent this, multiple labs have harnessed game engines and virtual reality (VR) [12, 13, 14, 15] to create immersive naturalistic tasks while still allowing for precise experimental control.

Our aim was not only to create naturalistic VR tasks in a way that would be easy to program and use, but to also make them comparable across multiple species. To meet these needs, we chose to create our VR tasks using Unreal Engine 4 (UE4) [www.unrealengine.com]. UE4 is a free state-of-theart game engine with open source code, with which realistic VR environments can be easily designed. Importantly, unlike other game engines such as Unity [www.unity.com], UE4 contains a mature visual scripting language that makes coding for non-programmers possible. Like all game engines, UE4 lacks features necessary for performing controlled behavioural experiments across multiple species, such as precise control over its timing. To implement these features in UE4 we developed DomeVR. DomeVR is a toolbox containing features necessary to create immersive and replicable behavioural tasks for cross species research and control them during execution.

In the following sections we will first introduce the features of UE4 used in our VR setup, and then we will outline the extensions provided by DomeVR.

### 2.1 Unreal Engine 4

In order to create DomeVR, we used multiple user friendly features present in UE4’s GUI. One extensive GUI (the Level Editor) is available to create “Levels” (or Maps) that are 3D areas the player (i.e. the experimental subject) can move through. Many aspects of the level’s Landscape can be modified; for example, through GUI tools like “Sculpt” or “Paint”, realistic levels with hills and mountains can be shaped. Levels are filled with Objects that can be drag-and-dropped into them. Objects that can be placed or spawned in the Level are called Actors. Actors can have Components attached to them which give them specific properties, for example how they are rendered or how they behave physically. Pawns are specific actors that can be controlled by Players (or AI) if they are possessed by a PlayerController (or AIController) and Characters are Pawns with the components for collision and movement attached by default. If a Character is possessed by a PlayerController, it becomes the character that is moved by the player’s input device (e.g. keyboard or joystick). Populating the VR environments with different Objects by simply dragging and dropping them into the Level makes UE4 an excellent choice for behavioural task design.

For Objects to interact in behavioural tasks, custom code must be added to them. UE4 provides a visual scripting language called “Blueprint”, which can be used inside the UE4 GUI and even enables non-programmers to add custom code to Objects easily. The same drag-and-drop principle is used to connect Blueprint “Nodes” like points on a flow-chart. Nodes can either be specialized UE4 class methods (e.g. “Get Game Time Since Creation” for Actors), control structures (e.g. “For Each” loops through an array), variables or literals. Using these elements, Blueprints programming interactions between Objects: e.g. code for an Actor to “Spawn” in front of the Player after a delay of a few seconds. Although Blueprint code is interpreted, it is compiled into fast byte code which offers a performance that is appropriate for real-time computer graphics. Interpreted code does not require lengthy compilation procedure and thus enables rapid testing of the written code. Due to the resulting much shorter coding cycles, we made use of Blueprints extensively and only used C++ for performance-critical parts or low-level code that would not be possible to code in Blueprints.

### 2.2 DomeVR

Our toolbox DomeVR has the following features:

1. A dome projection suitable for immersive human and non-human VR experiments.

2. Task control flow via state machines with blocks and trials.

3. Actors that can be placed in the virtual environment and serve as a part of tasks e.g. as a visual stimulus.

4. Actors for calibrating the visual field.

5. Real-time inputs and outputs e.g. eyetracking or eventmarkers.

6. Timing synchronisation to ensure data can be aligned to neural activity.

7. A logging system to automatically record relevant information needed for offline analysis.

8. Logging analysis in python.

9. An experiment GUI to control task parameters online and display behavioural performance.

The dome projection, eyetracking, eventmarkers and the log with timing synchronisation and analysis are available as independent modules that can be added into any UE4 project. You can find the code for the full DomeVR Toolbox and the extensible UE4 plugins at www.githubcom/zero-noise-lab.

We now describe the setup and the features of DomeVR one by one, after which their effectiveness and use is illustrated in a two-alternative forced choice task performed by mice, monkeys and humans in a dome environment suitable for all three species (see section 4).

## 3 Methods

The DomeVR toolbox is made up of many Blueprint or C++ code classes derived from UE4 base classes, the most important of which and their relationships between each other are outlined in Figure 1. These classes allow you to make any number of objects inheriting their class properties and methods. Their uses will be explored in the following sections.

**Figure 1:**
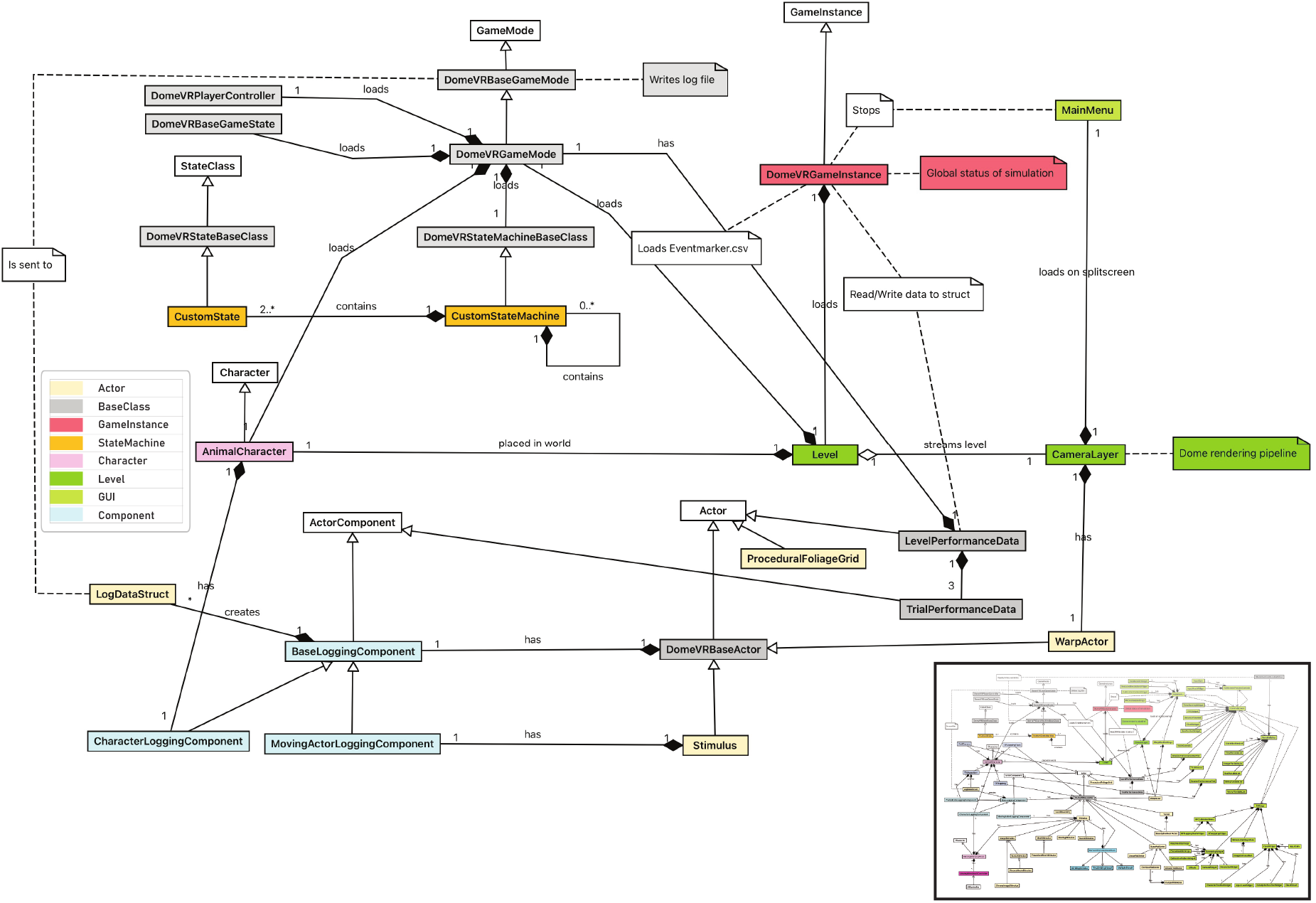
Overview of DomeVR. An overview of the major classes of the DomeVR project shown as colored nodes and their inheritance from UE4 base classes shown in white. The majority of classes are included in the inset, this is expanded on in Supplementary Figure 1. Note that not all symbols are Unified Modeling Language (UML) standard.

### 3.1 Dome projection

#### 3.1.1 VR visualization

To create an immersive virtual reality (VR) environment suitable for humans and non-humans we project the VR onto a curved dome. This creates an immersive experience for participants at the center of the dome by stimulating peripheral vision [16, 17]. For this purpose we used a Fibresports 60 cm radius spherical dome extending to 250^*°*^. The inner surface of the dome was illuminated using a 60Hz Canon XEED WUX450ST projector with a 1920×1200 resolution pointed at a 40 cm 180^*°*^ acrylic (PMMA) convex mirror (see Figure 2, for details of this method see Bourke [18]).

**Figure 2:**
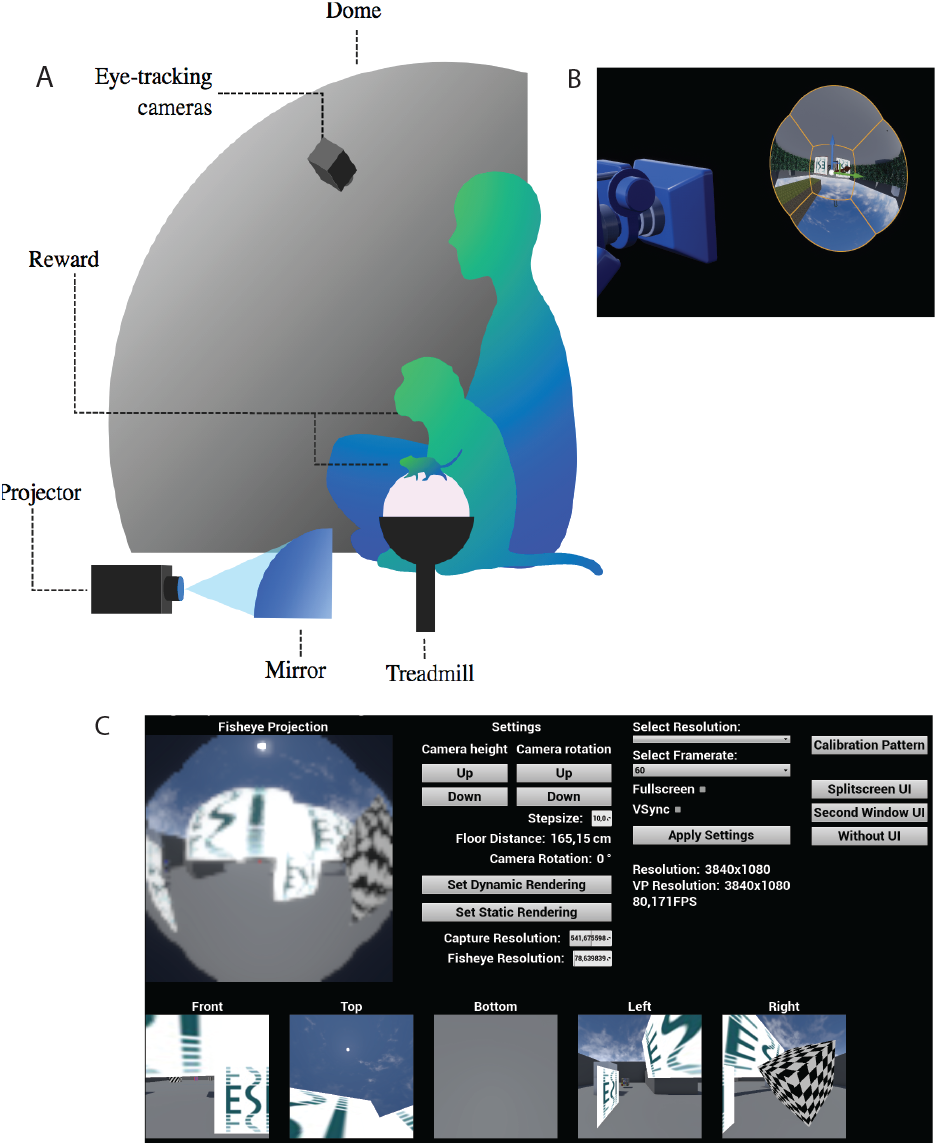
Dome projection (a) Schematic of the dome setup. The projector is pointed at the curved mirror that projects the UE4 output onto the 250 degree dome. Input is received through a treadmill/-trackball for mouse and primate participants respectively. A camera placed within the dome can track behavioural input e.g. eye move-C ments. (b) The fisheye view in Unreal Engine 4. The 6th scene capture component captures the warped fisheye view from the 5 different perspectives from 5 scene capture components. (c) The dynamic rendering pipeline UI allows the simultaneous viewing of the first 6 scene capture components and updating of their resolutions to find the best possible compromise between resolution and frame rate. Here demonstrated with a low resolution.

#### 3.1.2 UE4 dome projection

One of our major requirements was to create a spherical dome projection in UE4, which is necessary for an immersive VR environment that covers the full visual field of different species. In order to project a VR environmment onto a dome, the UE4 output needs to be warped. While a dome projection method for Unreal Engine has not previously been created, we were able to base our method on the method created by Paul Bourke for Unity [18] and therefore it is described here only briefly.

To create the dome projection multiple views of the virtual environment need to be warped together. Specifically, a (virtual) 5 camera rig of scene capture components was attached to the player and recorded the scene from the 5 different perspectives needed to cover a 250 degree dome. The scene captures were set to capture an image each frame and the capture sort priority was set to 0, which avoided tearing artifacts due to non-consistent rendering in the single scene captures. The captured image of each of these scene capture components was written to render targets with a resolution of 1024×1024, which provided a high enough sampling rate of the scene for a clear image in the final output. The render targets were then used as a texture source in a corresponding material and finally applied to meshes (downloaded from paulbourke.net) that distort the recorded images such that a fisheye view was created (see Figure 2B). The fisheye view was captured by a 6th scene capture component and the resulting image finally warped again by a mesh specifically created for our setup by the Meshmapper application of Paul Bourke. This step was performed for each setup individually. A WarpActor was created that allows for changing calibration meshes during run-time (see section 3.9). The distortion is captured by a final 7th scene capture component and rendered to the display. Since the position of the camera producing the output on the screen did not correspond to the position of the player, audio listeners for sound were manually attached to the players location. Additionally an 8th dummy camera was placed at the back of the character so that foliage would spawn automatically (see section 3.3).

While for the Unity method by Bourke [18] the cameras and meshes that create the dome projection could be hidden in an invisible layer, invisible layers that show only some meshes are not supported by UE4. Therefore, we first placed the assets for the projection outside the coordinates that we use to create our simulations (but close enough not to cause interpolation issues). Second, we set the materials to the unlit User Interface material domain to avoid any influence of reflections or lighting on our captured images. To create a uniform black background (used in some of our experiments), we further put them in front of a plane mesh with a black material. To reuse the render pipeline in other levels easily, we created the DomeRender level that only contains the described meshes for providing the fisheye projection and final dome distortion. Using the UE4 feature of Level Streaming, usually intended for splitting up the loading of large memory heavy environments, the DomeRender level (i.e. the rendering pipeline) was streamed to all levels.

We additionally created a dynamic rendering version of this pipeline to be able to test the settings of all the scene capture components in a UI (see Figure 2C). Using this UI, the correct resolution for the scene capture components can be chosen to reach a desired frame rate.

### 3.2 Control flow and states

A typical way to control the various stages of a task as well as the transitions between them (e.g. ‘start the trial’, followed by ‘let the stimulus appear’), is by using state machines (see Figure 3). State machines have a set number of States and move between them (e.g. from Start to Stimulus) with a Transition, which happens depending on whether a certain condition is true and/or an event dispatcher is called (e.g. GotoNext). We created a state machine using Logic Driver Pro - State Machine Blueprint Editor Version 2.4.6 (available to purchase here [19]) to control the flow of information during the task.

**Figure 3:**
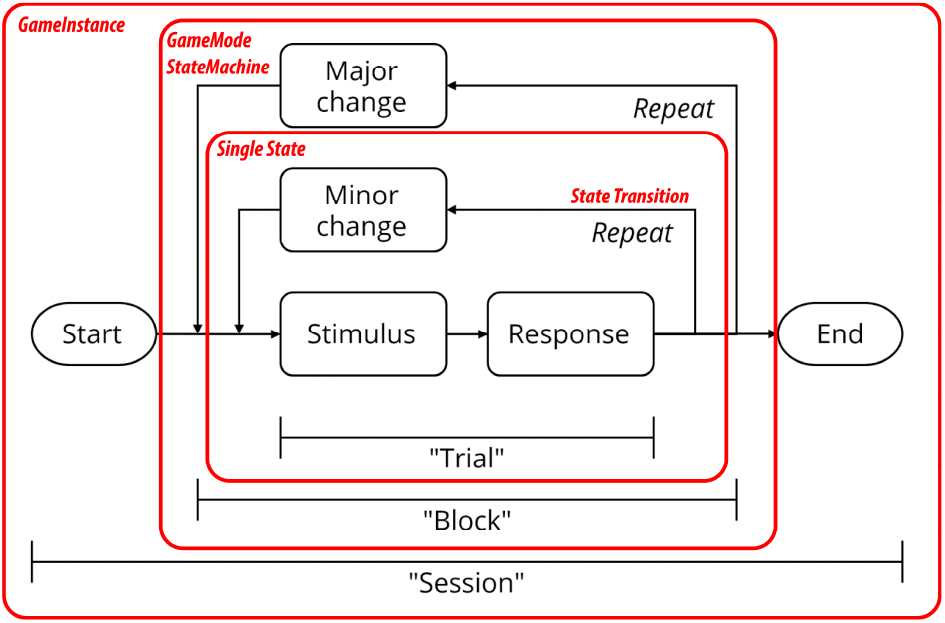
Graphic showing the stimulus display structure of DomeVR. A ‘Session’ describes the loading of a level for the subject to play. This loads a ‘Block’ state machine containing a ‘Trial’ state machine. Single states are shown as small boxes.

A “Session” is started by playing a Level with DomeVRGameInstance as a Blueprint which loads a State Machine. We took the approach of splitting these state machines into “Trials” and “Blocks”. A “Block State Machine” has at least one “Trial State Machine” as a State within it. The Block State Machine can define logic to create the correct version of the Trial, e.g. which stimulus should be displayed, as well as any other States necessary. A Trial State Machine consists of a collection of “Trial States” (described below). Both Trial and Block State Machines must inherit from the Blueprint parent class DomeVRStateMachine in order for the task events to be properly logged (see section 3.7). Visual blueprint scripting is very useful for the visualisation of the control flow within the state machines; see Figure 4 for an example Trial State Machine calling multiple Trial States. The flow of the code can also easily be visualised from the editor in debug mode. When nesting the Trial State Machines within the Block State Machine we used intermediate graphs. This is a functionality that allows the same Trial State Machine to be run with different parameters (i.e. to display different stimuli) such that the same Trial State Machine can be reused for different blocks of the experiment.

**Figure 4:**
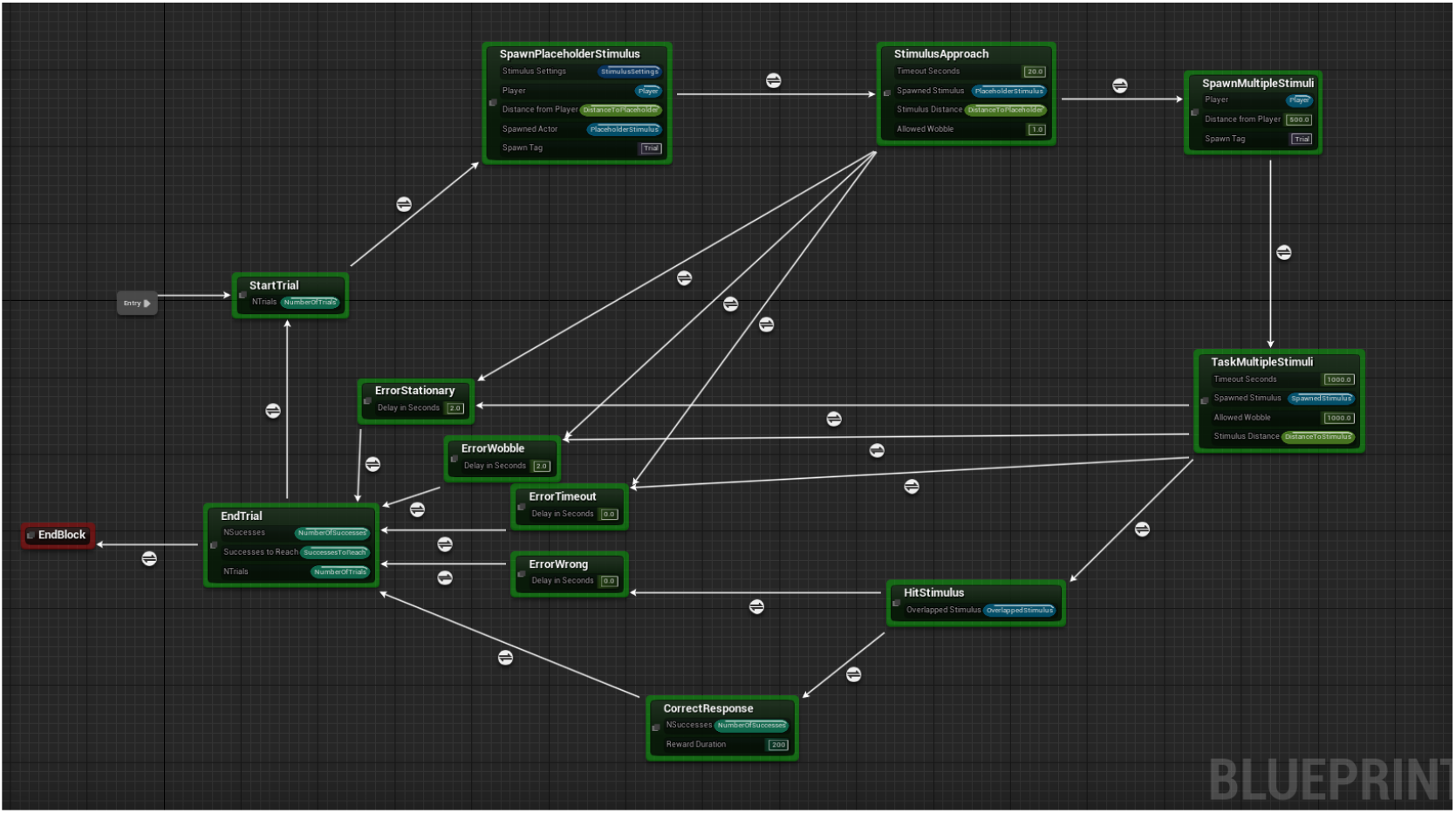
Blueprint code of an example trial State Machine. The grey entry box shows where the state machine begins and arrows connecting boxes show possible transitions between states.

Trial States contain the actual blueprint code that runs the experiment. They can have multiple input variables exposed on the node within the State Machine as shown in Figure 4; other inputs and outputs can be accessed from the node graph. Trial States were designed to be as simple as possible to aid their reuse across multiple State Machines. As for State Machines, Trial States must all inherit from a parent DomeVRBaseState class in order for their start and stop times to be tracked by the log (see section 3.7) and for other DomeVR specific features to be used (e.g. sending eventmarkers). To aid code readability we used certain standards, for example all TrialStates send a GoToNext event dispatcher when they are finished. Some example states are StartTrial, which increments an NTrials variable, and sends two eventmarkers: one that denotes the trial start and another to send the trial number (see figure 4). A more complex example is the SpawnMultipleStimulus state which takes in certain parameters and spawns a MultipleStimulus instance at the specified distance using the provided StimulusSettings struct (see section 3.4.1). It additionally sends multiple eventmarkers denoting these settings and parameters. Trial States are used as building blocks and reused across multiple StateMachines. They have their own unique Blueprint functions and events for control flow. For example, OnStateEnd is an Event which allows you to time Blueprint code to run only when the state is about to be exited. This system grants complete flexibility for the experimenter to create any type of task without needing to recode the entire environment each time.

### 3.3 Levels

To make realistic and immersive environments for our subjects, we made very large UE4 Levels (e.g. up to eight UE4 km long). For a realistic appearance of these levels, foliage (e.g. grass) needed to be placed abundantly while still maintaining a feasible performance at run-time. Therefore, we used UE4’s Landscape Grass Types, which provided the information for dynamic spawning of foliage. Materials were created that trigger foliage spawning as defined in the Landscape Grass Type, which was done dynamically around the camera position. If the spawned foliage exceeds the cull distance it was automatically deleted thus preserving performance even in large levels. To further save performance, the used meshes provided different levels of detail (LOD). An example of a ground material that used Landscape Grass types applied to the Landscape of a level is shown in Figure 5.

**Figure 5:**
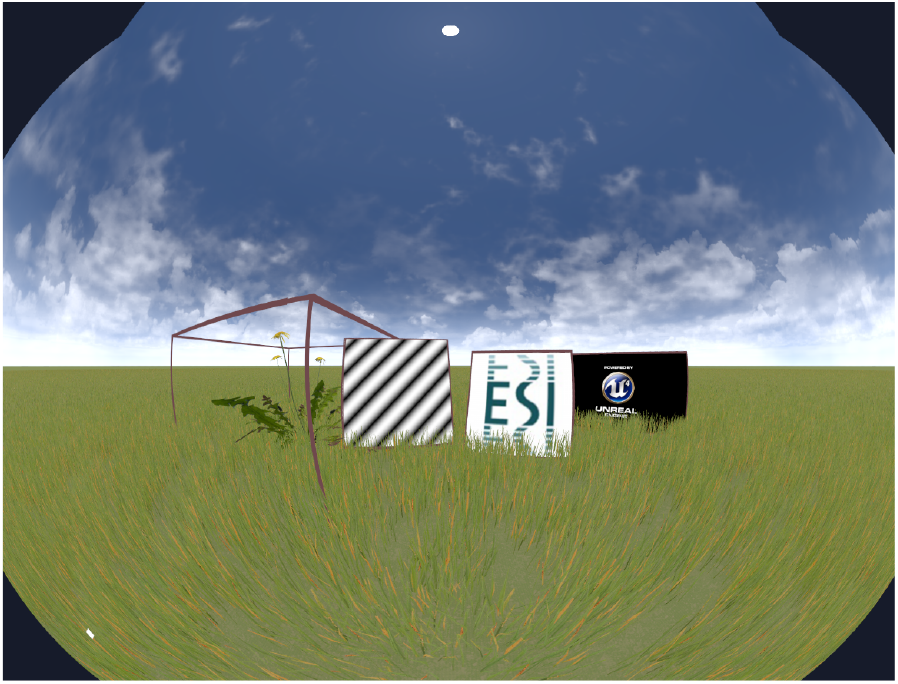
Examples of four Stimulus types added to the GrassyLandscape level. The grass was generated as outlined in section 3.3. From left to right: MeshStimulus, GratingStimulus, ImageStimulus, MovieStimulus

To be able to systematically determine how foliage is distributed throughout the level, we created a ProceduralFoliageGrid actor using the modified Procedural Foliage Placement Tool (PFT) (available to purchase here). This actor creates a grid that can randomly spawn foliage with a particular density that can vary between grid segments. The bounds of the actor fit the size of the landscape so that foliage can be spawned across the whole landscape. Labels show the chosen density within each grid segment. Once the foliage has been spawned it can be converted to one single Hierarchical Instanced Static Mesh (HISM) per foliage type so that it will be rendered efficiently and the level can be saved. The settings used to create the foliage are saved in a text file along with the random seed so that it can be recreated. Since this process is very memory and CPU intensive, it could only be performed on small landscapes for higher foliage densities. To avoid reaching the edge of the landscape, we additionally created a LevelBoundary actor which can be added to the level and automatically teleports the player back to the PlayerStart location if it nears the edge of the landscape.

### 3.4 Stimuli

#### 3.4.1 Experimental stimuli

In order to display stimuli (i.e. objects or images in the VR environment) from within UE4, we created a Stimulus base class to incorporate features that can be reused across Stimulus types. The Stimulus base class includes parameters like Scale (which changes the size of the stimulus), Height (which changes the height of the Stimulus from the ground) and Hide (which determines its visibility). Stimulus inherits from the DomeVRBaseActor so that it can log these parameters (see section 3.7). The Stimulus base class is inherited by our stimuli classes which include, among others, the stimulus types ImageStimulus, MovieStimulus, GratingStimulus, MeshStimulus (see Figure 5). Since Actor classes must be spawned or placed in the level for instantiation, Stimuli classes are accompanied by structs for their parameters, with one master struct called StimulusSettings. The master StimulusSettings struct defines the stimulus type to be spawned, contains the structs for each specific stimulus type and defines the basic parameters common to all stimuli. The StimulusType parameter switches between the type of Stimulus child class that should be spawned. This has the advantage that code does not have to specify which stimulus type it is expecting so one can spawn either an ImageStimulus or a GratingStimulus from the same State using StimulusSettings. Specific structs for each child Stimulus class allow their parameters to be easily set and applied to the StimulusSettings struct. These parameters are saved in the log as outlined in section 3.7.

The appearance of Actors in UE4 is determined by their Material. We created custom Materials via the Material Editor and added parameters to the materials that allow their properties, such as the emissiveness (e.g. glow) and opacity, to be changed. For example, we created an ImageStimulusMaterial for the ImageStimulus class which has a parameter ‘Image’ expecting a UE4 2D texture. This texture can be created from an image or supplied by an image path which loads it into a UE4 2D texture (using Rama’s VictoryBPLibraryfor loading images from hard drive or UE4’s Download Image node for web images). The ImageStimulus class then sets this ‘Image’ parameter from the StimulusSettings struct. A MediaPlayer texture is used similarly to display videos in the MovieStimulusMaterial for the MovieStimulus class. We additionally created Materials using Custom expressions. These materials then include HSLS code for shaders which allow us to display common stimulus types in neuroscience (e.g. gratings for GratingStimulus), with parameters (e.g. spatial frequency), that can be updated online.

Single and multiple mesh stimuli are shown using the MeshStimulus and ProceduralMeshStimulus classes. ProceduralMeshStimulus can use either UE4’s stock procedural mesh methods or the free RuntimeMeshLoader and free RuntimeMeshComponent to be able to load multiple self defined meshes. This allows the presentation of 3D stimuli (see Figure 5).

Since in many experiments two or more stimuli are simultaneously visible, a DerivedStimulusBaseClass was created so that stimuli can be spawned relative to a common root location. The MultipleStimulus class inherits from this and allows a flexible number of stimuli to be spawned at equal distances. It also creates an order for collisions so that only one is the overlapped stimulus (which we used as a decision parameter in the task). The TwoDividingStimuli class is similar but only spawns two stimuli and contains parameters that allows them to move away from each other in opposite directions.

#### 3.4.2 Receptive field mapping stimuli

Receptive field (RF) mapping stimuli are created in a different manner to the experimental stimuli outlined above. Since they are displayed relative to the dome coordinate system, they are direct children of the DomeVRBaseActor class. The three main RFmapping stimuli were RFmapping, a strip of a sphere that was traversed across the dome; RFMappingFlash, small gaussian blobs that can be flashed across the dome; and RFSparseNoise, black and white squares which can be displayed at random locations across the dome. All of these stimuli are attached to the player character so that they are in dome coordinates and have their own unique logging.

### 3.5 Input/output

In order for both human and non-human subjects to move through the virtual environment in a comparable manner we used modified mouse inputs via USB. For the primates a GK75-1602B 75mm trackball from NSI was used as input. Using a trackball as an interface with a computer is to our knowledge a novel method for macaques, but has been previously used by other non-human primates [20]. For the mice we used a 20 cm diameter Styrofoam ball suspended in the air (modified method from Harvey et al. [1]) with 2 Logitech G502 laser mice to read out its movement. A 3D printed holder for the ball was made according to open source schematics [21]. This was converted to an emulated Xbox controller as described in section 3.5.1.

#### 3.5.1 Ball input

While most common input devices (e.g. a joystick, lever or keyboard) can be used as movement input into UE4 natively, we have added the features necessary for trackball/multiple mouse inputs to be interpreted as movement (see Figure 2). We used UCR, a program which can be found on github, and plugins interception and ViGEm to emulate an Xbox controller and block mouse inputs from being seen by Windows and moving the cursor. The emulated Xbox controller was then selected and each axis of the controller bound to the correct InputAxis in the UE4 GUI (e.g. Forward Axis). InputAxis events of the Animal character were used to fine tune gain changes etc. for adjustment and online training purposes. The raw Input values were logged for analysis (see section 3.7 and example 2). Rotational input via Turn Axis was deactivated for experiments by removing the controller input for the Turn Axis in the Project Settings so that the player would always face in the same direction. Since the movement of the trackball or Styrofoam ball is continuous, we did not attempt to determine the exact timing of the mouse input relative to the raw Input values logged in UE4. However, based on previous tests of UE4 we expect some small variable delay in the order of 6 ms [22].

#### 3.5.2 Eventmarkers

National Instrument Data Acquisition cards are widely used to collect data with high precision. To facilitate the sending and receiving of precisely timed events and “eventmarker” codes we built an Unreal plugin to interface with DaqServer which can send and receive these events through very fast pipes. At the time of writing this supports cards: PCI-6221, PCIe-6321, PCI-6503, PCIe-6251, PCIe-6323, USB-6353 and PCIe-6351. In our setup PCIe-6321 was used. This program was designed specifically with neuroscience research in mind and has commands to provide reward of different lengths (TTL pulses on a specific card channel) and send an eventmarker code that can be used to signal the timing of particular events in DomeVR (e.g. the start of a new trial). We reserved the 16th bit as a strobe to control the timing of the reading of data and set this bit to high after each eventmarker is sent. Each time an eventmarker is successfully sent, a Query Performance Counter timestamp is noted (for more details see section 3.6).

#### 3.5.3 Eye tracking

Eye tracking can be performed by interfacing with EyeServer. We built an Unreal plugin UnrealEyeserverInterface to interface with the Eye Server, which supports iRecHS2 [23] and Eyelink (SR research). These are both camera based eye tracking systems which are able to output the subject’s calibrated eye position at high frequency in real-time. Thus far we have exclusively tested iRecHS2. UnrealEyeserverInterface sends requests to iRecServer to query the current eye position, request that certain eye windows are tracked, and can query whether or not the eye is within the window. Eventmarkers can be sent to iRecHS2 via a UDP connection in order for the timing of the saved data to be synchronised with the DomeVR log. To calibrate the eye the plugin can accept points in the iRecHS2 window so that UE4 can be used to display the calibration points.

### 3.6 Timing control

As UE4 is designed for smooth gameplay, it does not natively have the millisecond timing precision necessary for neuroscientific experiments. It instead uses multiple threads (“GameThread”, “RenderThread”, “RHI Thread”) which are all processed and displayed to the screen as quickly as possible, such that there is no consistent delay between the time that a frame is finished being processed in the GameThread and the time that it is displayed on screen by the GPU. To account for these variable delays, we made use of the DirectX IDXGISwapChain used by UE4. DirectX IDXGISwapChain is a collection of already rendered frames that are waiting to be displayed on the screen. Using its GetFrameStatistics method, we can access the SyncQPCTime, which gives the Windows QueryPerformanceCounter timestamp of the most recent screen flip, the index PresentCount and the UE4 GFrameCounter gives the index of the current GameThread frame in the swap chain. By additionally recording the QueryPerformanceCounter timestamp of each eventmarker we were able to adjust the timings to align with when they appeared on our dome. All this information is recorded in our log as ViewportFrameStatistics. These timing statistics can only be accessed when the game is both fullscreen and compiled (i.e. not when played in an editor window).

In order to check that these modified timings were correct we used a photodiode to record light level changes on our dome (see section 4.1). A Flickerpattern actor attached to the player switches between brightness or colors on each frame when a state machine is running. We attached this actor to the player to ensure that it was always in the same location on the dome. A widget in the GUI (see section 3.9) allows the user to move it to the correct position. This Flickerpattern actor also sends eventmarkers at each color switch so that the timing of each frame can be checked. We recorded the brightness changes of the Flickerpattern using an amplified photodiode that was built in-house. Both the photodiode signal and the 16 bit eventmarkers were recorded at 30 kHz using an Open Ephys acquisition board. This Open Ephys system was modified in-house to accept 16 channels of TTL inputs. The timing statistics were saved via DomeVRLog (see section 3.7).

### 3.7 Behavioral logging

In order to reconstruct what happens during a DomeVR task we created a customized logging system which we call the DomeVRLog. This is defined in the DomeVRBaseGameMode C++ class and implemented by the BaseLoggingComponent attached to children of the DomeVRBaseActor class (e.g. Stimuli).

Two UTF-8 encoded plain text files are automatically created when a new level is loaded (see Section 3.3). One contains all information (hereafter referred to as the “continuous log”) and one contains only some of this information for convenience (hereafter referred to as the “behavioural log”). The header of both logs contains many parameters about the experiment as a whole, including the level start time, Subject, Experiment, etc. (see Listing 1). Each line of the file after the header contains a minimum of 4 columns. First is a column with the UE4 game thread time of the logged event, second is a column with an “Object identifier” (which is a unique number that can be used to track the identity of all spawned objects in the UE4 world) and a column specifying the “LogTypes” of the line, for example SpawnLocation. The final columns depend on which LogType the line is. Since this log is saved in plain text it is human readable and therefore future proof. In order to efficiently parse this large text file we created a python module using memory mapping that is open source (see section 3.8).

**Listing 1:**
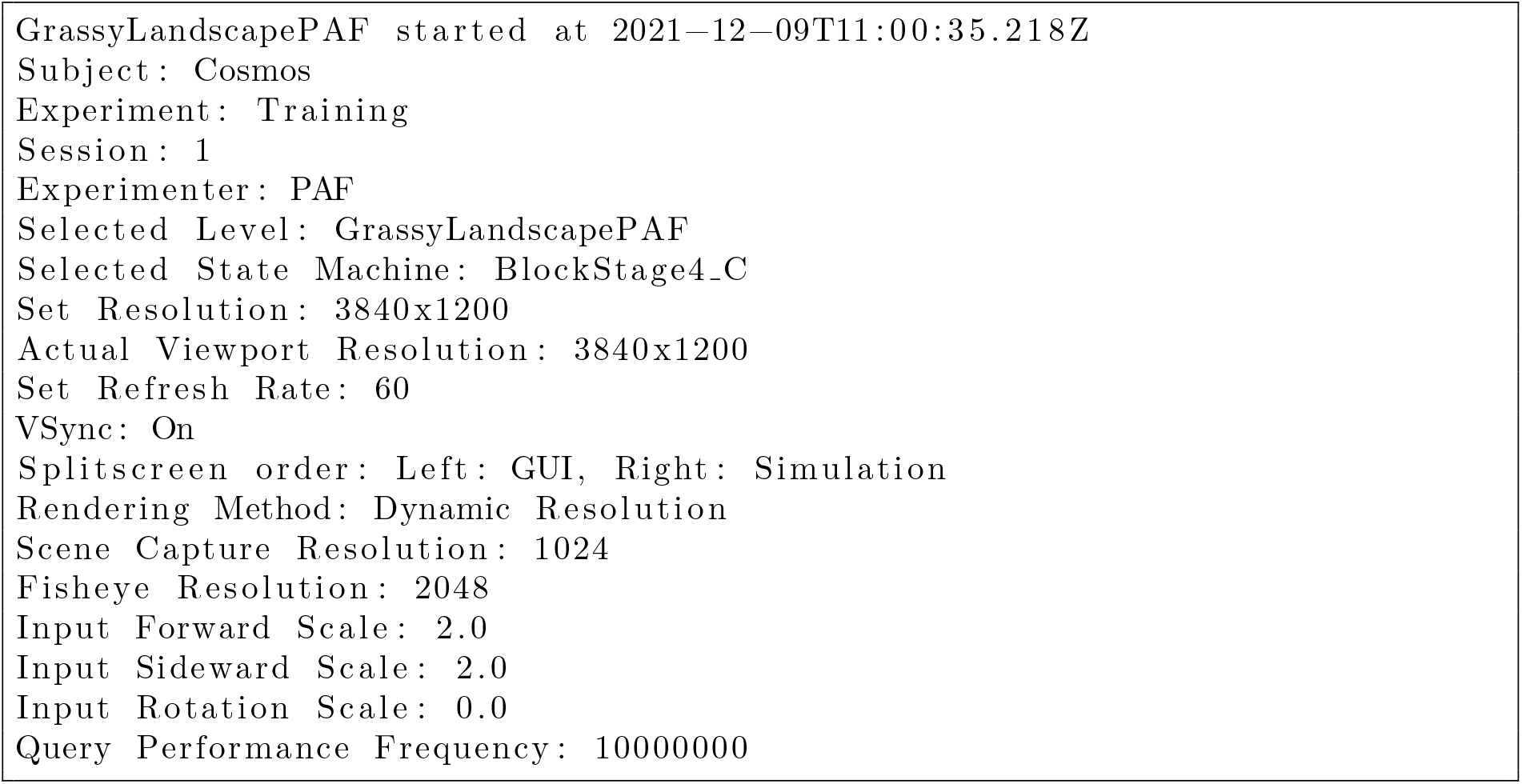
Example log header

The amount of information contained in the behavioural log is determined by setting a verbosity level. At the lowest verbosity level only the spawned actors, destroyed actors, specific log messages and state changes are logged. Increasing the verbosity level gradually increases the amount logged until it approaches the continuous log. This log can be used for behavioural data analysis.

The continuous log is meant to be an account of everything that happened during the experiment. For example, it contains the spawn and destroy point of every DomeVR stimulus shown during the session, stimulus parameters on initialization or on update and the entry and exit of all DomeVR states (see section 4 and Listing 2). Additionally, on each tick information about the player is written by the CharacterLoggingComponent, for example, the Cartesian Unreal coordinates of the player in the world. Also, with each tick the position of all actors with a MovingActorLoggingComponent (e.g. stimuli created from the DomeVR classes) are logged relative to the player in both dome (the radial position of the bottom of the stimulus in the dome) and Cartesian coordinates (in Unreal units). Sending of 16 bit eventmarkers are automatically logged (see section 3.5), and there is also the possibility to send custom information. The timing of the log is based on the time of the UE4 game thread using the GetRealTimeSeconds function. However, the actual display time is delayed relative to this due to the additional time taken to render the scene. From the 3rd game tick onwards the ViewportFrameStatistics are logged to adjust the GameThread time to the actual screen time (see section 3.6).

**Listing 2:**
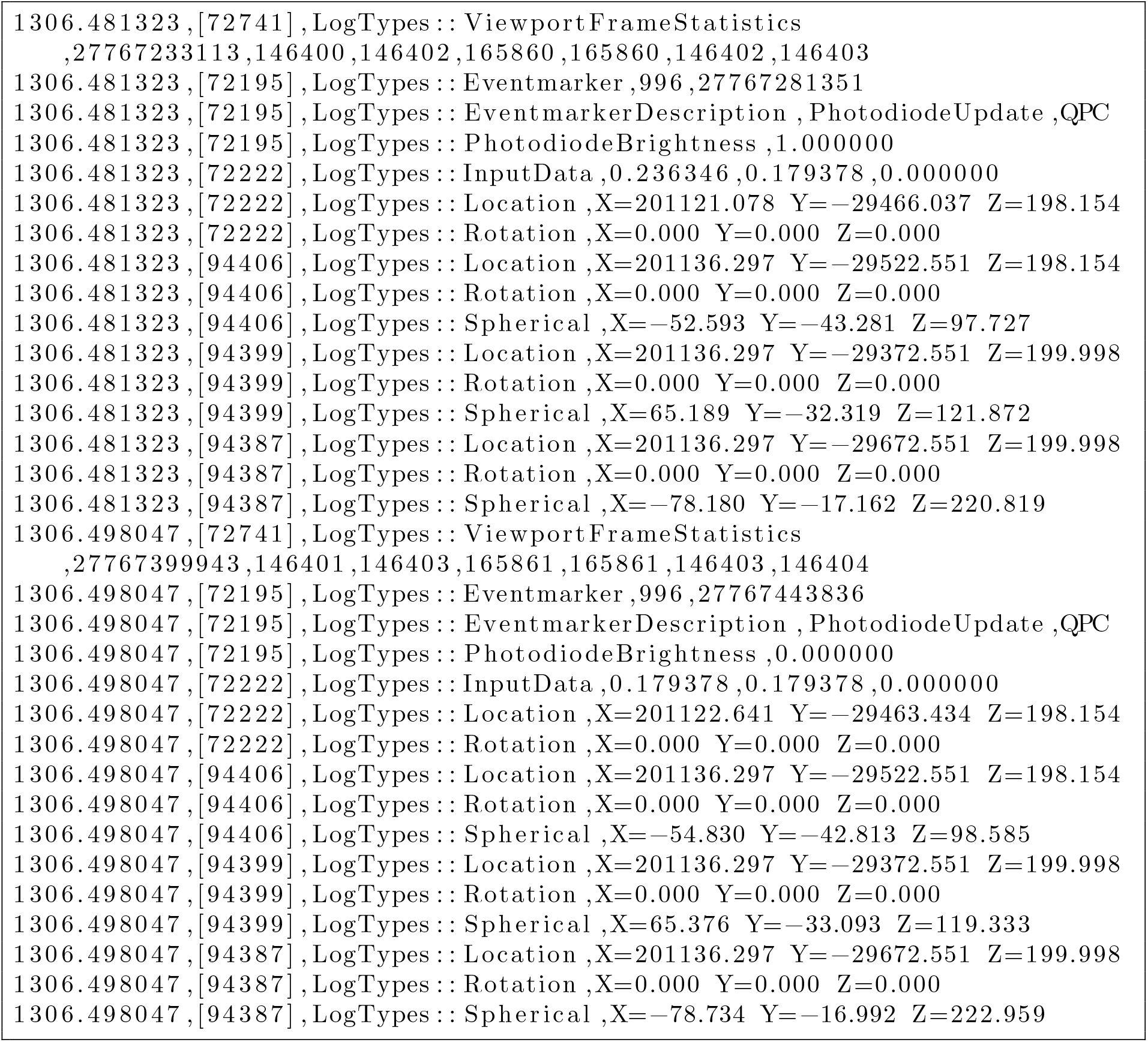
A two tick long excerpt of an example continuous log

### 3.8 Log analysis

In order to analyse the data stored in the DomeVRLog we created a parsing module parse domevrlog using Python. Since using States and StateMachines means that the outline of a trial can be completely flexible we needed the DomeVRLog to be parsable in a similarly flexible manner. We handled the large file size of this human readable text format by making use of memory mapping and on-demand loading to allow the DomeVRLog to be parsed quickly. The file index of the spawning time of every UE4 object (AnimalCharacter, StateMachines, Stimuli) was automatically stored for reference. In addition, automatic adjustment of all log timings to screen time were performed (see Section 3.6), according to the measured frame rate (59.952 Hz), the number of steps in the rendering pipeline (2) and projector delay (18 ms) (e.g. the delay from the time of the screen flip to the time it was displayed on the dome) was also taken into account. This was all done automatically so the user received the adjusted timestamps. Finding the location of the player between two states (in this case StartTrial and EndTrial) was done through flexible functions like `parse all state times and parse position`, see the following example:

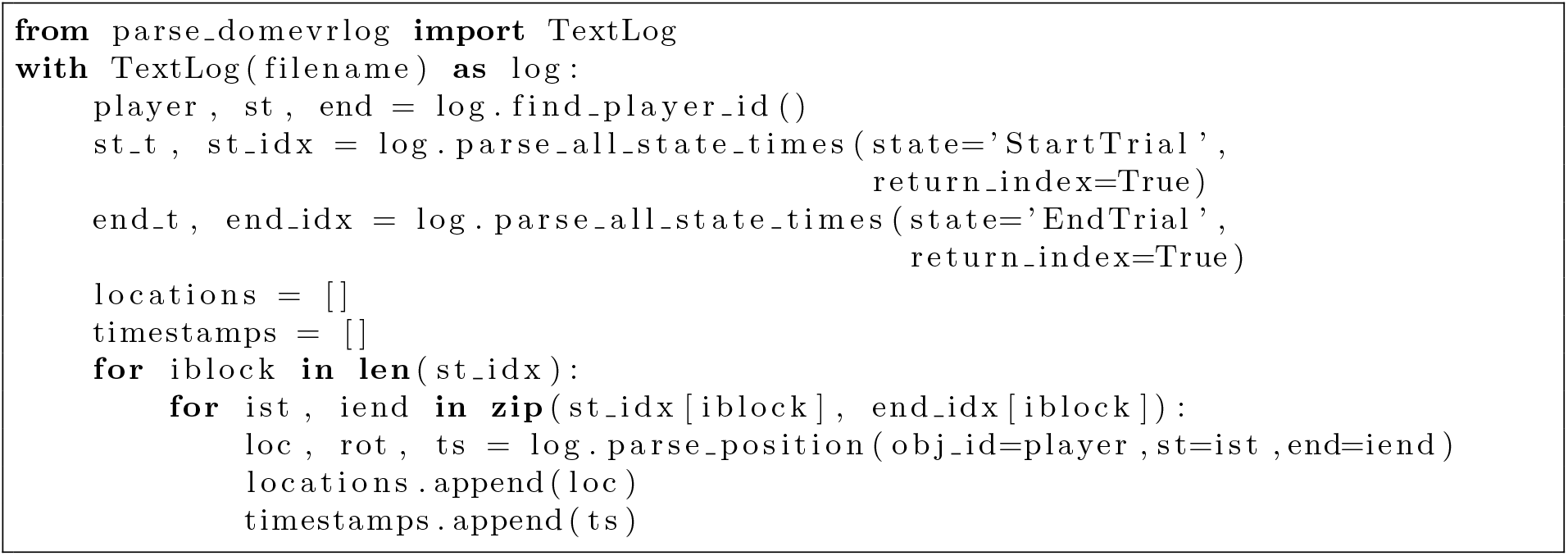

### 3.9 Graphical user interface (GUI)

In a final step we created a GUI to help scientists flexibly change their experimental settings, view the performance of the subject and debug issues with their experiment. We created this GUI using the Unreal Motion Graphics UI Designer (UMG). It was displayed on a second “experimenter” screen while the dome projection was displayed on the projector. The first part of the GUI, that starts automatically when DomeVR is run, is MainMenu. This contains fundamental settings for starting a task, including selecting the Level and the State Machine that will be run, as well as saving the subject and experimenter name (see Figure 6A). These settings must be selected here or they are not logged. State Machines and Levels stored in the appropriate folders are automatically detected and available to select from a drop down list. Settings from the previous session are stored in the Saved folder of the project. On the first run for a new user a pop-up requires the user to select the dome warp mesh (see section 3.1.2), photodiode settings (see section 3.6) and input adjustments (see section 3.5). When the RUN button is pressed the selected State Machine and Level are run (as well as the NidaqServer and EyeServer if requested) and the GUI switches to ControlScreen.

**Figure 6:**
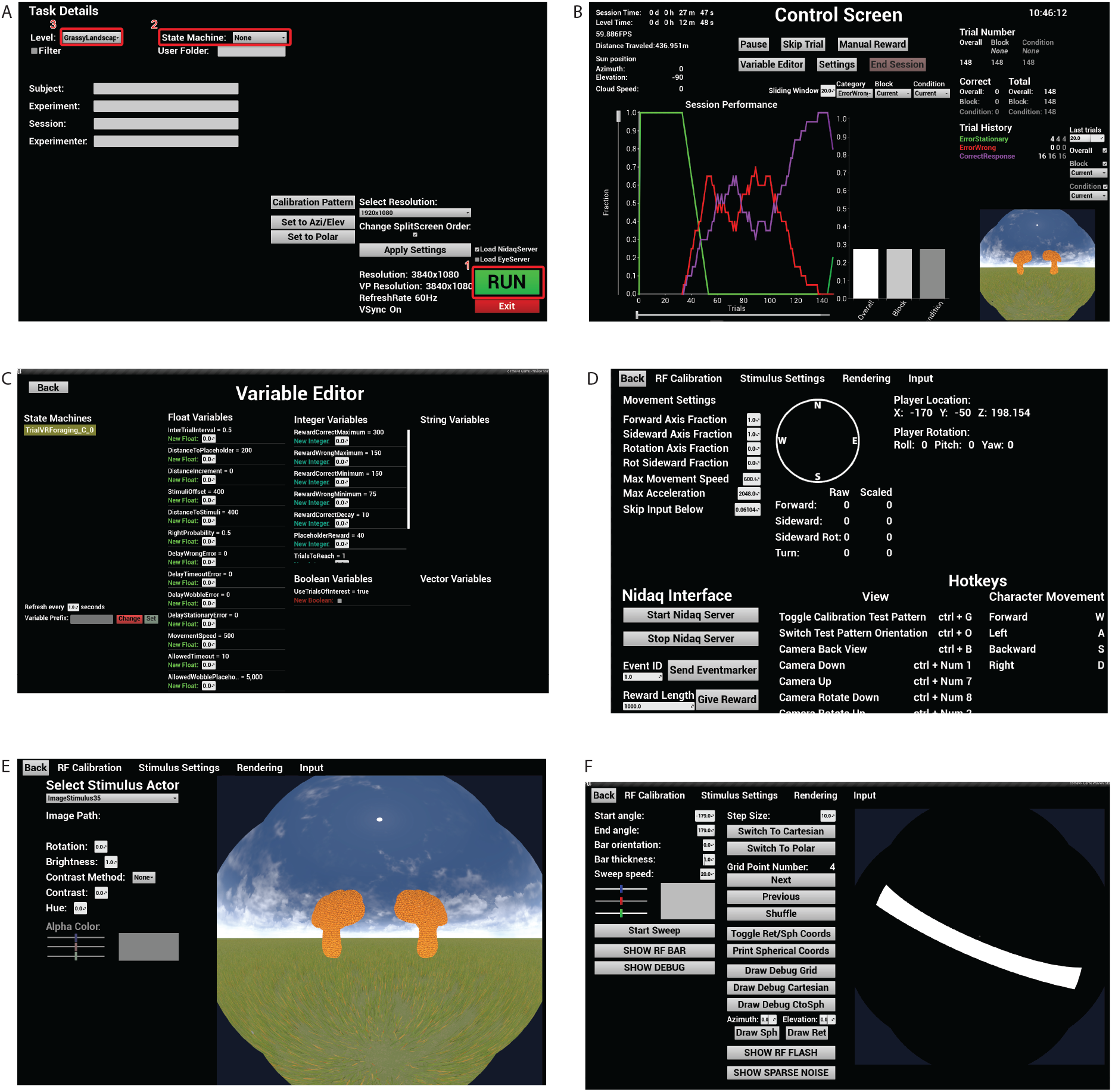
Different tabs of the GUI. (a) The Menu has 3 necessary buttons to run the task: (1) Button to start running with the selected settings. (2) Drop down menu to find StateMachines in the Blueprints folder. (3)

The ControlScreen GUI contains buttons for interacting with the running experiment as well as displaying information about the performance of the subject and information about the current UE4 Level (see Figure 6B). A bar chart displays the average performance per condition and a line chart displays the time series of performance per condition across the last n trials using the free Kantan Charts plugin (available from the Unreal store). The ControlScreen also displays an image of the FisheyeProjection (see Figure 2B) in the bottom right corner so that the experimenter can view the subject screen. Clicking on it opens a larger view of the fisheye projection where input settings can be changed. Other interactive components of the ControlScreen include a “Pause” button (which also blanks the participant screen out), the “Variable Editor” (see section 3.9.1 and Figure 6C), and the “Settings” button which brings up tabs of other widgets, for example an Input tab (see Figure 6D) to see the vector angle of the participant etc. and a Stimulus Settings tab (see Figure 6E) which allow the selection of a spawned stimulus and editing of its StimulusSettings properties online. These were used for debugging issues with the experimental setup.

Drop down menu to find Levels in the Level folder. (b-e) The various other tabs of the GUI that are present when a task is run. (f) The GUI for receptive field mapping parameters.

To allow users to click on the ControlScreen for the interactive components while simultaneously maintaining timing control of the subject screen as outlined in section 3.6, we needed to maintain fullscreen focus on the DomeVR window across two screens. To achieve this we used NvidiaSurround to bind the dome projector screen to the experimenter screen creating a virtual single screen with a resolution of 3840×1200. The warped dome projection was on one half and the GUI on the other half and the screen refresh time of both are synchronised. This is a major advantage over many other stimulus presentation software in which only keyboard shortcuts can be used to change settings during timed stimulus presentation.

#### 3.9.1 Variable Editor

The VariableEditor widget displays variables sorted by their type (e.g float, integer) from the running state machines and allows them to be changed online (see Figure 6C). Since any variable can be changed with this widget, a filter is applied by default such that only variable names starting with EDIT are displayed. The VariableEditor interface relies on a back-end C++ class called StateMachineVariableEditor. It uses the UE4 reflection system to read out variable names and values to exposed TMaps, which are used to build up the variable lists in the GUI. For setting the variables to a different value, the StateMachineVariableEditor class provides exposed functions that are called when a value is changed and set in the GUI, respectively. When two state machines are active (e.g. a block StateMachine that calls a trial StateMachine) the variables from both can be accessed by selecting the corresponding state machine from the list of active state machines in the VariableEditor. To update the displayed variables, a new StateMachineVariableEditor instance from the selected state machine is created and all variables are refreshed as explained before.

#### 3.9.2 Performance Counters

To display the performance of the subject on the ControlScreen (see Figure 6B) their performance in each category was stored in counters. These counters were not predefined but were created (or incremented) from any state during run-time by calling IncrementCounter with a counter name. Maintaining all of these counters was done by the LevelPerformanceData actor. The counting was divided into three scopes referring to trials, conditions and blocks. For each of these scopes, separate counters were maintained, which was done in three TrialPerformanceData actor components attached to the LevelPerformanceData actor. The scope could also be switched by any state. In addition to the performance charts, the values of the counters were shown in the trial history at the right side of the ControlScreen GUI, which could filter for different blocks and conditions as well as display the counted value for a certain number of trials back in time.

### 3.10 Example experiments

We tested DomeVR on subjects using Unreal Engine 4.24 on custom built Windows 10 PCs from Alternate with 32 GB RAM, 1 TB SSD, AMD Ryzen 9 CPU and either a Geforce RTX 3090 or Nvidia Titan RTX GPU. The human experiments were performed with permission from the ethics committee of the Medical Faculty of Goethe University (No:2021-252); the mouse and monkey experiments were approved by the Regierungspräsidium Darmstadt (No:F149/2000).

Here we present a task built with DomeVR that was performed by three different species. This task was inspired by Havenith et al. [15], and is the subject of another manuscript (Crider et al, in preparation) and is therefore described only briefly here. All three species were asked to distinguish two natural shapes, embedded in a grassy field in a simple, two-alternative choice task. We present example data from 1 human, 1 monkey and 1 mouse. The shapes for the humans and monkey varied smoothly between a starfish and a flower shape (see Figure 9A); the mice had to distinguish between a jagged and a round leaf. On each trial, a blend between these two ‘extreme’ stimuli was shown alongside a reference stimulus (the middle blend between the two extremes). Mice only needed to distinguish between extremes. The target stimulus (the shape more resembling the starfish and the jagged leaf respectively) was rewarded with a click sound for the humans, a drop of juice for the monkey, and a drop of vanilla soy milk for the mice; the distractor stimulus did not yield any reward and for mice it was accompanied by white noise and a time-out. The humans and monkey indicated their choice by moving to the stimulus with the 75mm trackball; the mice ran towards the stimulus on the floating styrofoam ball (described in section 3.5).

## 4 Results

### 4.1 Photodiode timing measurements

The timings of the eventmarkers relative to the photodiode signal were adjusted by the log parsing script according to the QPC timestamps recorded in the log (see section 3.6 for details). To show that the eventmarkers were correctly adjusted we plotted the recorded photodiode signal from the time that it was turned on in a session. In order to identify the correct frames we created a Flickerpattern with four steps (see Figure 7A). The adjusted photodiode timing and level clearly matches the recorded signal.

**Figure 7:**
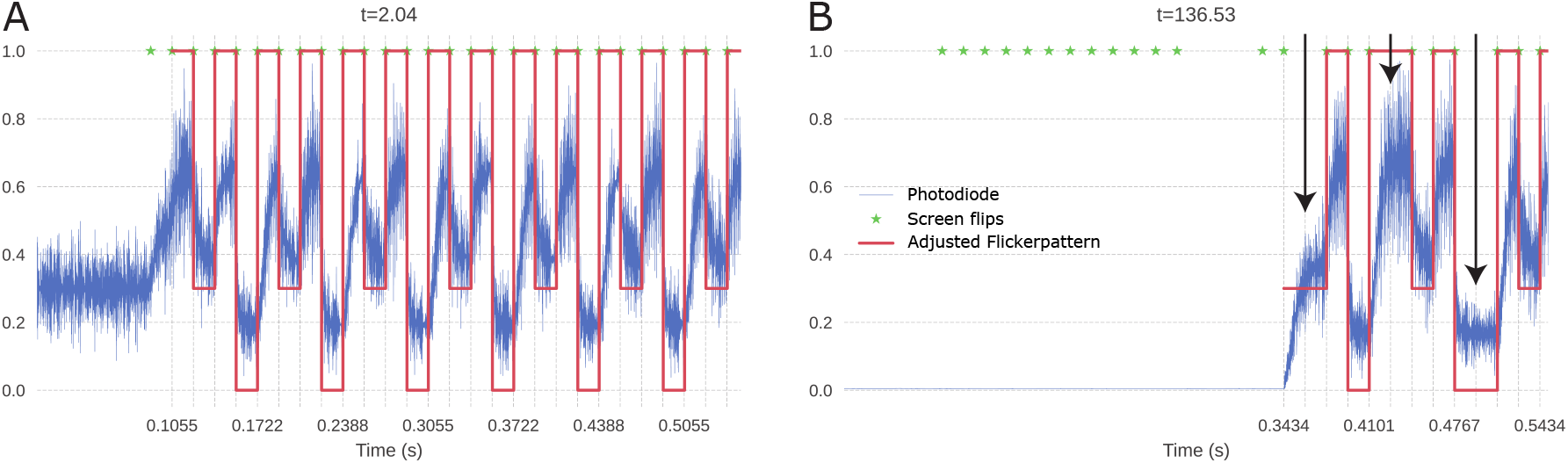
Photodiode results (a) Photodiode recording of the Flickerpattern with three brightness levels in the following pattern: 1,0.3,1,0. It initializes with a brightness of 0.5. Blue line is the recorded photodiode signal, green stars are all screen flips and red line is the software adjusted Flickerpattern. (b) Photodiode recording of the Flickerpattern after a pause, meant to induce frameskips (see arrows).

To ensure that our timing adjustments were correct even in the event of frame skips (stuck frames that are shown for twice the normal length of time) we caused them to happen by pausing and restarting the game. During a pause the whole screen was set to black. In Figure 7B three such frame skips can be seen indicated with arrows and the timings from the log follow them exactly.

To measure the accuracy of the adjustment we aligned the photodiode traces of the Flickerpattern switching from a brightness of 1 to 0 at the time indicated by the IDXGISwapChain screen time alignment. In Figure 8A we see all traces in the session including frame skips (n = 41,904 transitions). To quantify the accuracy of this alignment we measured the time of the maximum signal change. This resulted in a cluster of points around 0 ms with a voltage of above 1 (see Figure 8C). The median time of the timing distribution was −0.6667 ms, with the 5th and 95th percentile at −0.7333 ms and −0.5667 ms respectively. The fact that the flip appears to begin before time 0 suggests that our measured projector delay of 18 ms was in fact too long for these highly accurate alignments and would ideally need to be measured to μs accuracy. Therefore, even in this session of enforced frame skips, 90% of the traces were aligned to within 0.1667 ms.

**Figure 8:**
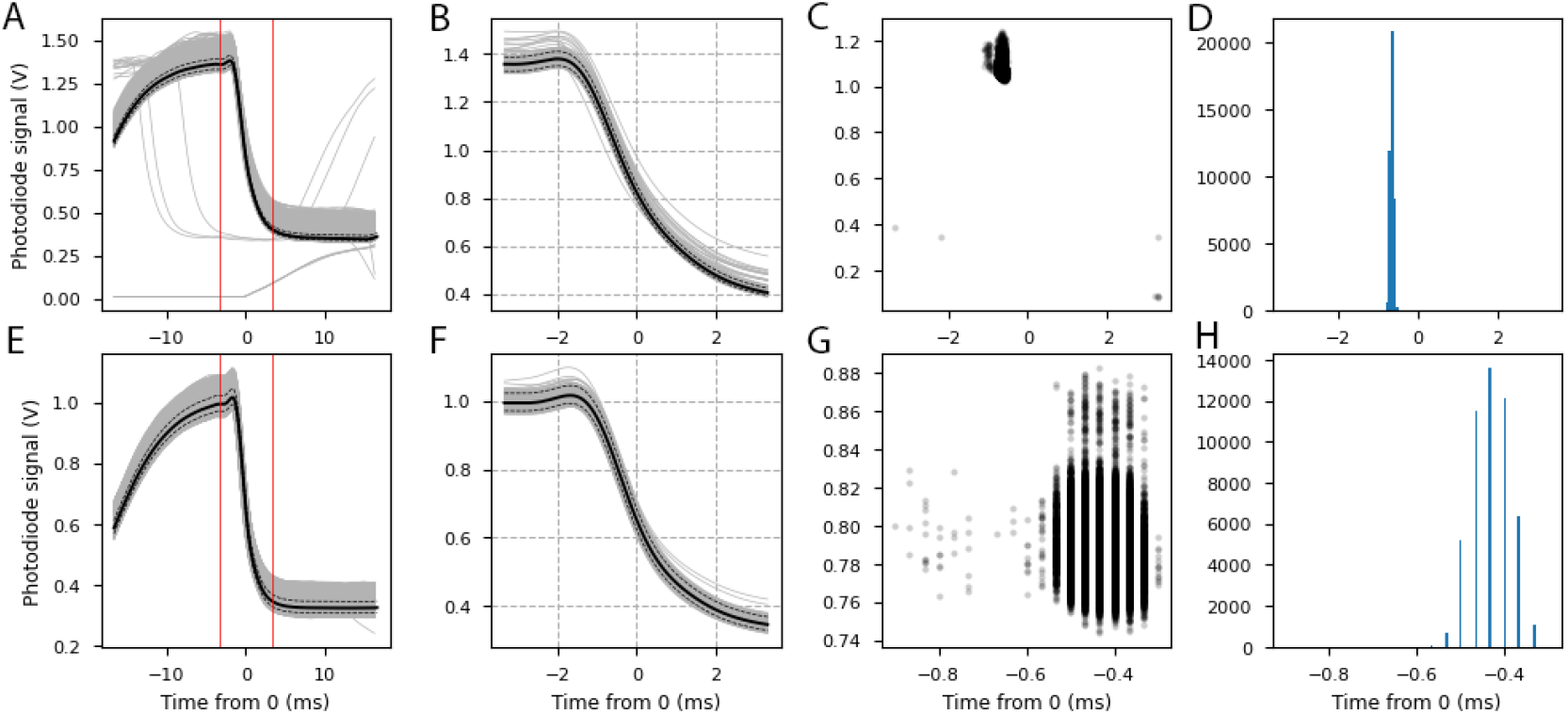
Photodiode traces aligned by IDXGISwapChain timings. (a) Analysis of photodiode alignment. Grey lines are individual alignments, black line is the median, dotted lines are 5th and 95th percentile. Red lines indicate analysed time window. (b) As a but for the shorter analysed time. Note the change in x axis. (c) Black dots are the maximum change in the photodiode signal of the analysed area. (d) Histogram shows the counts of the maximum change times. (e-h) As a-d but the session was run on a second computer and DomeVR was not paused. Note the change in x axis in g and h. Here the maximum change varies by less than a millisecond and the limit of the 30 kHz recording resolution of 0.0333 ms is visible.

For comparison, we repeated the analysis using a session on a second computer without pauses or frame skips. This time all Flickerpattern transitions from 1 to 0 are well aligned (n=50656, see Figure 8E-H). Therefore the time of maximum signal change is even more tightly clustered. The median was −0.4333 ms with the 5th and 95th percentile at −0.5 ms and −0.3667 ms respectively. In this ideal session 90% of the traces were aligned to within 0.1333 ms and in both ideal and non-ideal sessions 98% of the traces were aligned to within 0.2 ms.

### 4.2 Example case experiment

To demonstrate the utility of our setup to create tasks across species, here we show data from three example sessions across three species; human, macaque and mouse. We used similar Levels containing open grassy landscapes in all species but the StateMachines are slightly different to comply with the needs of each species (e.g. reward and punishments were not necessary for the human subjects). The details of the task are outlined in section 3.10.

In Figure 9 we see a screenshot from a human subject performing the task. In Figure 9A on the left is the experimenter screen with ControlScreen GUI and Fisheye view of the task during the session. On the right is the warped view projected onto the dome that the participant saw. In Figure 9B-D the paths of the subjects for the first 50 trials in Unreal coordinates are shown. These paths demonstrate that all three species are able to accurately navigate in the virtual environment. Figure 9C demonstrates that a macaque was able to control a trackball with a high degree of precision. Finally 9E shows the performance at the conclusion of the human, monkey, and mouse sessions.

**Figure 9:**
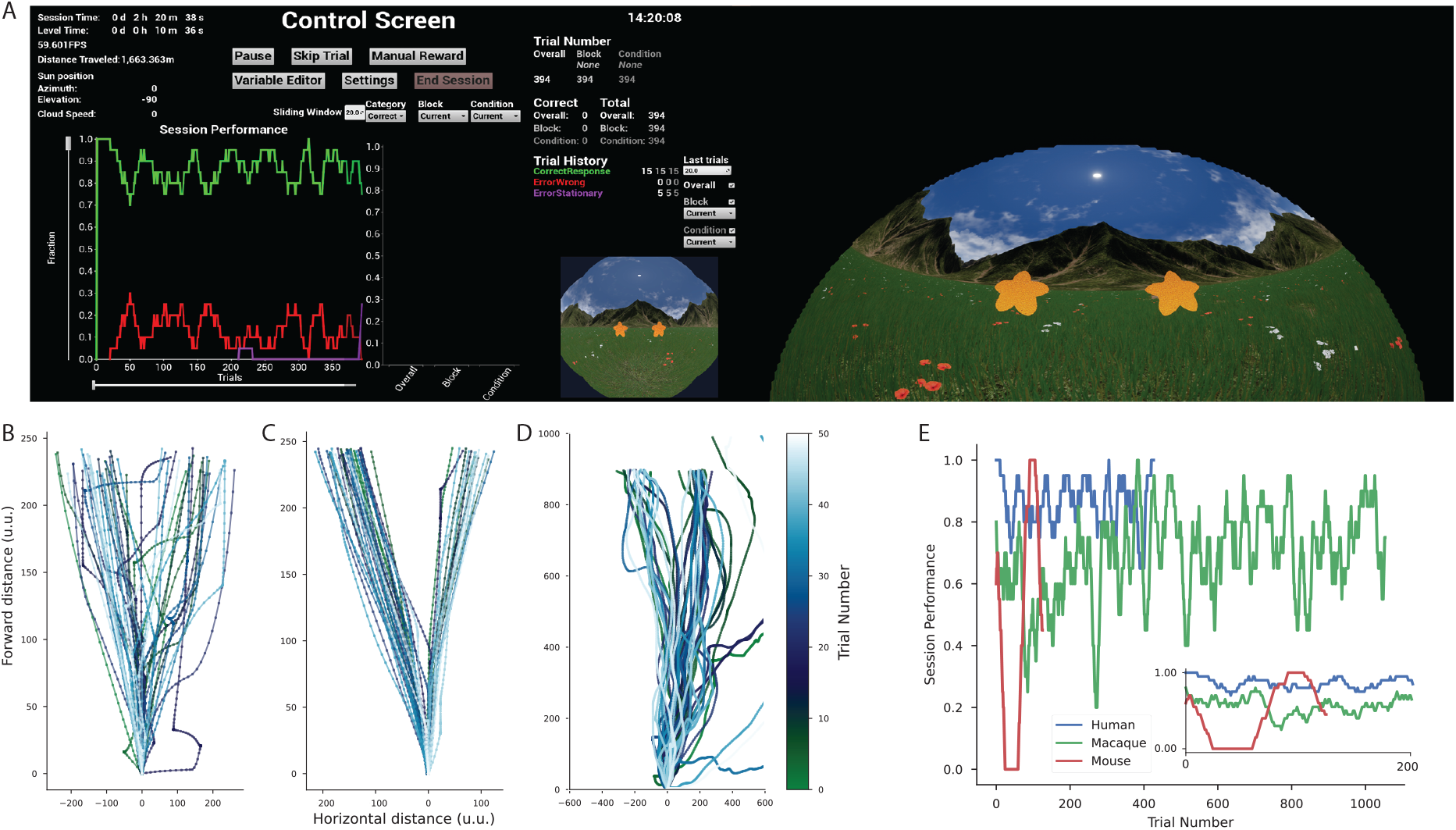
Example data from three species, human, macaque and mouse. (a) Dual screen screenshot of a human experiment (b) The path to the stimuli from the first 50 trials of a human. (c-d) As b but for the macaque and mouse experiments respectively. Different x and y axis in each are due to the different parameters of each experiment. (e) Comparative performance across trials for all three species. Performance is measured as the proportion of correct trials in a sliding window of 20 trials. For the human this is the same as the green line in a. Inset shows the same performance measure for just the first 200 trials of each experiment.

## 5 Discussion

Here we present a versatile, easy-to-use toolbox for creating VR environments, making use of the powerful game engine Unreal Engine 4. Arguably, UE4 is the best game engine in use for easily generating photo-realistic VR environments, with drag-and-drop features for creating detailed scenes and builtin naturalistic lighting, physics and more. We coupled this to the flexible experimental control and high-precision timing needed for neuroscientific experiments. Our DomeVR toolbox has three crucial advantages. First, many components of it are modular and can therefore be used independently in other UE4 projects. Second, our toolbox can be used across several species by allowing different types of inputs, such as an air-suspended running ball for mice and eye tracking for monkeys, and by projecting the VR in a dome that covers the visual fields of all typical model species. This is crucial to create a true, immersive VR experience in species with laterally positioned eyes such as rodents. Third, our toolbox allows users with little to no programming experience to create experiments via the use of

Blueprints. Blueprints are essentially a visual interface that transforms code into interconnected graphs, which represent typical game elements such as variables and events. They effectively give the user the speed and power of programming in C++, without ever being in contact with this conceptually difficult programming language. Other commonly used VR game engines such as Unity and Panda3D, generally require proficiency in C# or Python respectively.

### 5.1 Comparison with other VR toolboxes

Over the years, numerous toolboxes have been created to use VR in neuroscience experiments. Many of these have been developed specifically for investigating spatial navigation in humans (e.g. PyEPL[24], MazeSuite[25], PandaEPL[26], EVE[27], VREX[28], UXF[29], NavWell [30], bmITUX[31], Landmarks[32] and OpenMaze [33]). Almost without exception, these toolboxes use Unity to create their VR environments. On the other hand, VR experiments for mice are usually written directly in graphical programming languages such as Blender [4, 15, 34] and Bonsai [14] and typically feature simplistic, non-naturalistic VR environments. The only VR toolboxes for primates that we are aware of are the VR toolbox by Doucet et al. [12]; and the Unified Suite for Experiments (USE; [13]), which is tested on primates, humans, and AI.

The VR toolbox by Doucet et al. [12] is the only other VR toolbox for neuroscientists that is written with Unreal engine. It runs on Unreal Engine 3, the previous version of UE4, which does not feature Blueprints but instead uses the visual scripting language Kismet. Kismet has a much reduced functionality compared to Blueprints. To make the toolbox as accessible as possible, it is controlled through text commands sent via TCP, which can be sent using any programming language. It therefore uses two computers, one for running Unreal and the other for controlling Unreal using these TCP text commands.

This has the disadvantage that information sent via these predefined TCP text commands is limited compared to the many options available in the DomeVR GUI. The TCP connection also provides a hurdle for precise synchronization, which the authors overcome by synchronizing the computer clocks every 16 s through the Automachron software and logging the resulting time difference, which was 0 ± 3 ms. However, the accuracy of this synchronisation was not tested for the timing of the events on the screen, which we synchronised offline in DomeVR. The toolbox supports any input device that can be configured as a mouse (e.g. a joystick or a trackball), and could therefore in principle be used for primates and rodents – though it has so far only been tested on a flat screen for humans and monkeys.

The other VR toolbox for non-human primates, the USE toolbox [13], is written in Unity. It offers excellent timing precision through the use of a piece of external hardware, the USE SyncBox. This SyncBox recieves the timing of a photodiode and game engine ticks in order to synchronise them offline. Our synchronisation method is software based and therefore does not require a secondary piece of hardware to perform the timing precision while still achieving sub millisecond precision. Apart from this, the USE toolbox is very comparable to ours in what it achieves, with our toolbox offering two important advantages: the projection into a dome, which makes it suitable for rodent research, and the use of Blueprints, which makes it possible for non-experienced programmers to nevertheless code experiments in our toolbox.

A promising toolbox for rodent VR research is BonVision [14]. BonVision is an open-source software based on the Bonsai visual programming language, which can be used for displaying virtual reality (e.g. in a dome) as well as standard (2D) visual stimuli (e.g. on a flat screen). Its timing accuracy is in the order of 2 frames. The package contains several basic 3D objects such as spheres and cubes, and can import more complex natural scenes created in other VR programmes such as Blender. However, it is unlikely to be able to achieve the photo-realistic immersive tasks we have created. Through its use of Bonsai it is able to integrate with commonly used neuroscientific hardware natively and in a closed-loop manner. Similarly to Blueprints, it is suitable for non-programmers.

### 5.2 DomeVR issues and extensions

In the future, the DomeVR toolbox could be further developed to make it more generic. For instance, Task States could be added for a more diverse task structure than the focus of our laboratory on perceptual decision tasks featuring two simultaneously presented stimuli. Specifically, features for spatial navigation tasks, such as the easy creation of mazes, might be added, since VR is most commonly used in this domain. The toolbox could also be adapted to use on a flat screen instead of a dome. Finally, the added ability to measure delays in the processing of input by UE4 would be useful for many labs. We have provided open-source modules for eye tracking, events and eventmarker sending, dome projection and logging with timing synchronisation. Unfortunately, other parts of the toolbox had to be based on paid-for UE4 extensions, which are therefore not freely accessible. We believe if more neuroscientists were to embrace Unreal Engine, the dual goals of photo-realism and ease of use for non-programmers could be met in an open source manner.

## 6 Conclusion

In summary, we have created a VR toolbox that provides a vastly superior and more naturalistic environment than any of the rodent VR software around, and is the only one that creates a truly immersive VR experience in primates. It features flexible experimental control and accomplishes extremely precise experimental timing. It allows non-programmers to code their task designs flexibly via the use of Blueprints, and many components of it are modular and can therefore be used in other UE4 projects. Although this toolbox was created primarily for use in monkeys and mice, it is versatile enough to be used in humans also, and in fact we have tested it on those three species, with very comparable results. We are excited to witness a general transition in systems neuroscience away from static task designs towards more naturalistic, active tasks, and hope that DomeVR will contribute to this transition by providing other neuroscientists with the means to easily implement such tasks.

## Acknowledgements

We would like to thank Pierre-Antoine Ferracci, Robert Taylor and Anna-Katharina Cramer for data collection. Additionally we would like to thank Joscha Schmiedt, Georg Haas and Michael Stephan for technical support.

This work was supported by funding from the Max Planck Foundation (MPFS).

## Notes

### Competing Interest Statement

The authors have declared no competing interest.

