## Supplementary Figure for "DomeVR: A setup for experimental control of an immersive dome virtual environment created with Unreal Engine 4"

### Supplementary Figures

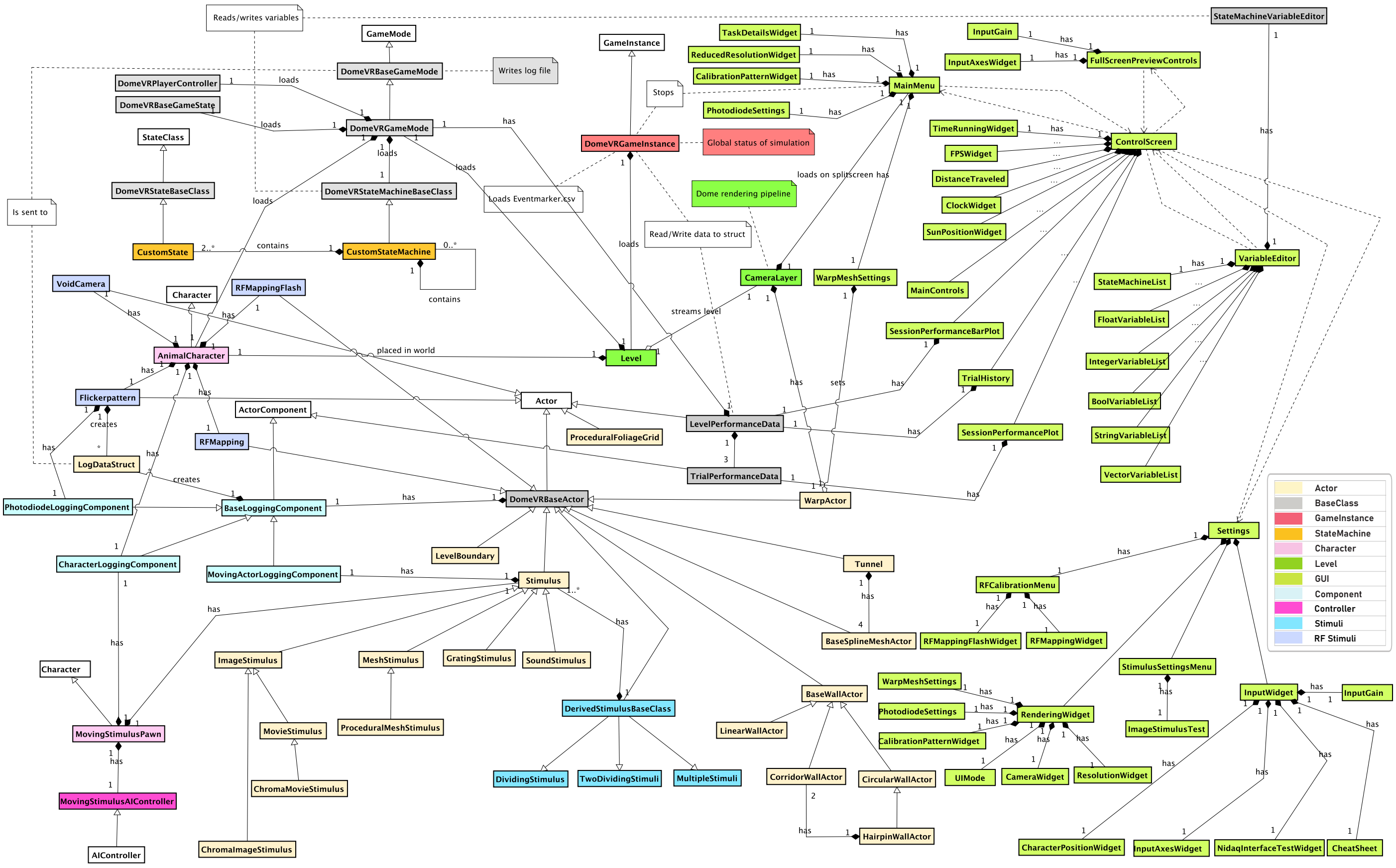

Supplementary Figure 1: DomeVR classes. An overview of most classes of the DomeVR project shown as colored nodes and their inheritance from UE4 base classes shown in white. Note that not all symbols are Unified Modeling Language (UML) standard.
